# BMPR-2 gates activity-dependent stabilization of dendrites during mitral cell remodeling

**DOI:** 10.1101/2020.10.30.358861

**Authors:** Shuhei Aihara, Satoshi Fujimoto, Richi Sakaguchi, Takeshi Imai

## Abstract

Developing neurons initially form excessive neurites and then remodel them based on molecular cues and neuronal activity. Developing mitral cells in the olfactory bulb initially extend multiple primary dendrites. They then stabilize single primary dendrites, while eliminating others. However, the mechanisms underlying the selective dendrite remodeling remain elusive. Using CRISPR/Cas9-based knockout screening combined with *in utero* electroporation, we identified BMPR-2 as a key regulator for the selective dendrite stabilization. *Bmpr2* knockout and its rescue experiments show that BMPR-2 inhibits LIMK without ligands and thereby facilitates dendrite destabilization. In contrast, the overexpression of antagonists and agonists indicate that ligand-bound BMPR-2 stabilizes dendrites, most likely by releasing LIMK. Using genetic and FRET imaging experiments, we also demonstrate that free LIMK is activated by NMDARs via Rac1, facilitating dendrite stabilization through F-actin formation. Thus, the selective stabilization of mitral cell dendrites is ensured by concomitant inputs of BMP ligands and neuronal activity.

## INTRODUCTION

In the mammalian nervous system, functional neuronal circuits are established via a circuit remodeling process during early postnatal development. Neurons initially form excessive neurites. Later on, however, they strengthen some neurites, while eliminating others. For example, at the neuromuscular junction and the cerebellar climbing fiber–Purkinje cell synapses, multiple axons compete for one target. While each synaptic target is initially innervated by multiple axons, only one establishes strong synapses, and all the other connections are eliminated (Lichtman and Colman, 2000; Watanabe and Kano, 2011). In the barrel cortex, Layer 4 neurons preferentially orient their dendrites toward thalamocortical axonal inputs through selective dendrite remodeling (Iwasato and Erzurumlu, 2018). It has been assumed that the neurite remodeling is a result of interplay between molecular guidance and activity-dependent processes (Valnegri et al., 2015; Wong and Ghosh, 2002); however, we do not fully understand how these factors orchestrate neurite remodeling. A long-standing question in the field is how some neurites are “selectively” strengthened and others are weakened during the remodeling process.

To study the mechanisms of selective neurite remodeling, mitral cells in the mouse olfactory bulb (OB) are an excellent model system. In the olfactory system, olfactory sensory neurons (OSNs) expressing the same type of odorant receptor converge their axons onto a set of glomeruli in the OB. OSN inputs are then relayed to the second-order neurons, mitral and tufted cells, in the glomeruli of the OB. A single glomerulus is typically innervated by 20-50 mitral/tufted cells (Imai, 2014). Each of the mitral/tufted cells connects its primary dendrite to a single glomerulus. Early in development, mitral cells extend multiple dendrites to multiple glomeruli; however, they stabilize some, but destabilize other dendrites over time. By the end of the first postnatal week they have eliminated all but one “winner” primary dendrite. Eventually, the winner primary dendrite forms thick tufted structure within a glomerulus (Blanchart et al., 2006; Fujimoto et al., 2019; Lin et al., 2000; Malun and Brunjes, 1996). Our recent study demonstrated that spontaneous neuronal activity in the OB is required for the pruning of supernumerary dendrites (Fujimoto et al., 2019). However, activity does not eliminate all the dendrites. It has remained unclear how a particular primary dendrite in a neuron is selectively stabilized during the activity-dependent remodeling process.

In this study, we screened for cell surface receptors that control dendrite remodeling process in mitral cells, employing CRISPR/Cas9-based knockout vectors combined with *in utero* electroporation. We found that BMPR-2 plays a critical role in selective dendrite stabilization during the remodeling process. In the absence of ligands, BMPR-2 inhibits LIMK1 through its intracellular tail domain, which causes dendrite destabilization. On the other hand, ligand-bound BMPR-2 promotes dendrite stabilization, most likely by releasing LIMK1. Free LIMK is activated by neuronal activity via NMDAR-Rac1-PAK pathway, resulting in F-actin formation and dendrite stabilization. Thus, dendrites are selectively stabilized when BMP ligands and glutamatergic inputs co-exist.

## RESULTS

### Screening for cell surface receptors regulating dendrite remodeling in mitral cells

To gain insight into the mechanisms of selective dendrite remodeling, we screened for cell surface receptors that control dendritic growth and/or remodeling based on extrinsic cues. Firstly, we obtained transcriptome data for developing mitral cells labeled by *in utero* electroporation. Single-cell cDNA was prepared from mitral cells at P3 and P6 and gene expression profiles were analyzed using a microarray (Imai et al., 2009) (See **Methods** for details). We focused on cell surface receptors abundantly expressed in developing mitral cells (**Figure S1**).

To test their functional roles in dendrite development, we performed CRISPR/Cas9-based knockout (KO) screening combined with *in utero* electroporation **(Figure 1A)**(Straub et al., 2014). To maximize the KO efficiency of any gene, we used the Triple-Target CRISPR method: Three different guide RNAs (gRNAs) were designed per gene and introduced to neurons together **(Figure S1A)**(Sunagawa et al., 2016). We electroporated tdTomato, Cas9, and the three gRNA plasmids into mitral cell progenitors at E12 in order to label and manipulate gene functions in mitral cells. It is unrealistic to examine the KO efficiency for all genes, as antibodies are not available for all of them. However, control experiments with two representative genes, *Tbr2* and *Tbx21*, demonstrated a highly efficient KO ratio with this strategy: 73% and 68% of tdTomato-positive mitral cells showed a complete lack of protein expression, respectively (**Figure S1B-E**). We therefore performed *in vivo* KO screening for the candidate genes (some were double or triple KO during the initial round of screening). OB samples were collected at P1 and/or P6, and dendrite morphology of mitral cells was comprehensively analyzed by using tissue clearing, SeeDB2, and volumetric imaging **(Figure 1A, B)**. Among the 38 genes tested, *Bmpr2* demonstrated the most prominent defects in dendrite development **(Figure S1F, G)**. When analyzed at P6, 93% of wild-type mitral cells formed just single primary dendrites. However, only 64% of mitral cells formed a single primary dendrite when *Bmpr2* was knocked out; the remaining neurons retained connections to multiple glomeruli **(Figure 1D and S2A)**. We did not find any obvious differences between the control and *Bmpr2* KO at P1, excluding a role for BMPR-2 during the initial dendritic outgrowth **(Figure 1C)**. Thus, BMPR-2 is either directly or indirectly involved in the dendrite pruning that occurs during the remodeling process.

**Figure 1.**
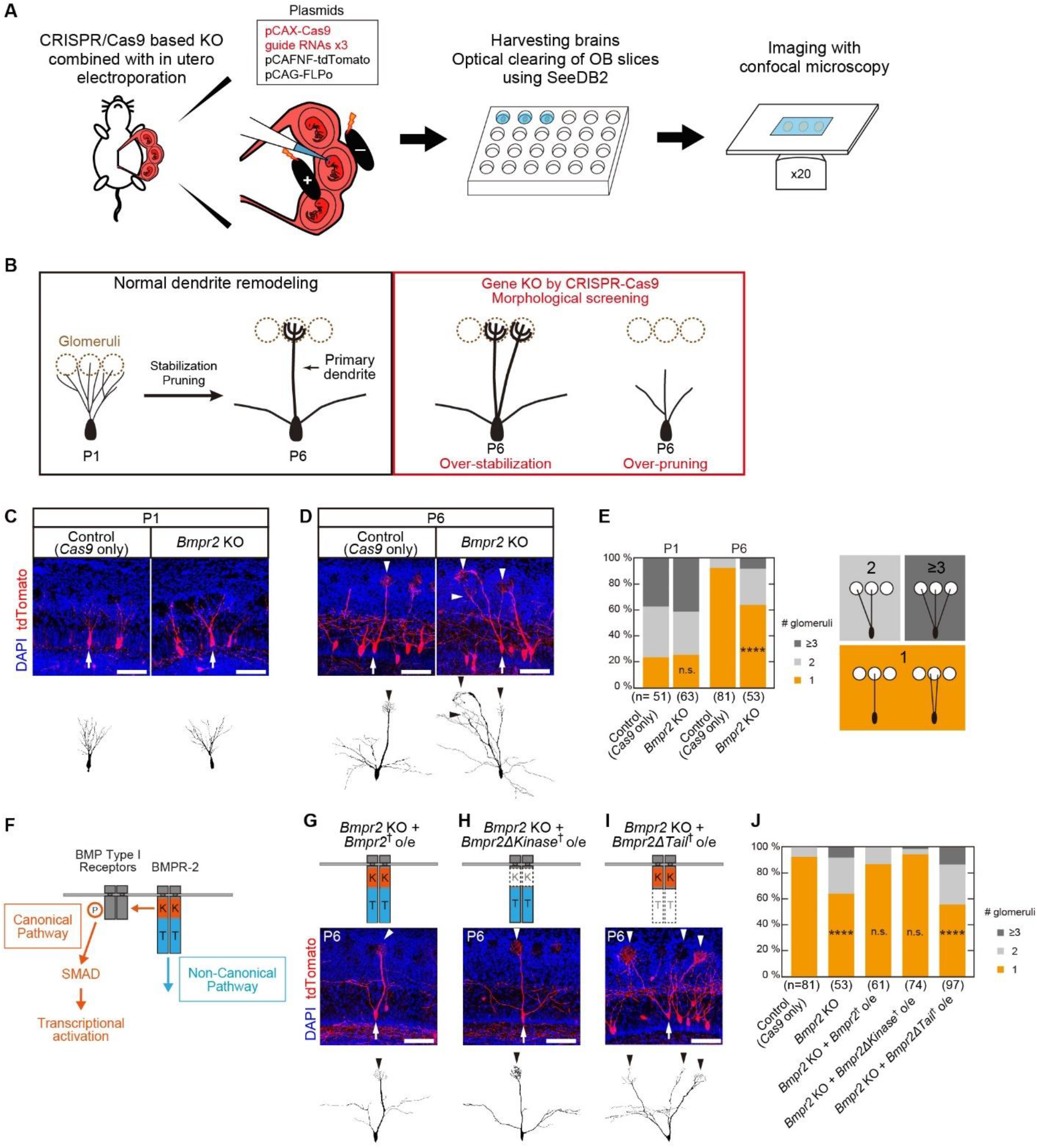
BMPR-2 tail domain is required for dendrite destabilization in mitral cells. **(A)** A schema showing CRISPR/Cas9-based KO screening. We designed three gRNAs for different exons for each gene. Plasmids for Cas9, the gRNAs, tdTomato, and FLPo were introduced by *in utero* electroporation at E12. Fixed brains were sliced, cleared by SeeDB2, and imaged with confocal microscopy. **(B)** Development of mitral cells from P1 to P6. A mitral cell establishes a single primary dendrite through the dendrite stabilization and pruning processes. We screened for genes controlling dendrite remodeling based on the morphology of primary dendrites at P6. **(C, D)** Representative images of control and *Bmpr2* KO mitral cells at P1 (C) and P6 (D). Arrows and arrowheads indicate somata and primary dendrites, respectively. Representative neurons are reconstructed and shown below. Scale bars, 100 μm. **(E)** Quantification of the number of glomeruli innervated per mitral cell. n.s., non-significant. **** p<0.0001 (χ^2^ test, compared to the control). n, number of mitral cells. **(F)** A schematic representation of the two BMP signaling pathways. The canonical pathway is mediated by a kinase domain of BMPR-2, BMP Type I receptors, and SMADs, leading to transcriptional activation. The non-canonical pathway is mediated by the C-terminal tail domain of BMPR-2. K, kinase domain; T, tail domain. **(G-I)** Rescue of *Bmpr2* KO phenotype with deletion mutants of *Bmpr2*. Guide RNA-resistant *Bmpr2* was used for the rescue experiments. † indicates a guide RNA-resistant *Bmpr2*. Age, P6. Arrows and arrowheads indicate somata and primary dendrites, respectively. Scale bars,100 μm. **(J)** The number of glomeruli innervated per mitral cell was quantified. The C-terminal tail domain is essential for dendrite pruning in mitral cells. n, number of mitral cells. n.s., non-significant. **** p<0.0001 (χ^2^ test with Bonferroni correction, compared to the control).

### The BMPR-2 tail domain is required for the normal dendrite remodeling

BMP signaling plays important roles in various aspects of development, from embryonic body patterning to neurite and synapse formation (Bragdon et al., 2011; Dutko and Mullins, 2011). BMP receptors are comprised two types, Type I and II. BMPR-2 is a member of the Type II receptor group. Their intracellular signals are divided into canonical and non-canonical pathways **(Figure 1F)**. The canonical pathway is initiated by the phosphorylation of the Type I receptor by the kinase domain of the Type II receptor, leading to transcriptional activation through SMAD proteins. The non-canonical pathway is mostly mediated by the C-terminal tail domain of BMPR-2 receptors (Foletta et al., 2003). To determine which pathway is required for normal dendrite remodeling in mitral cells, we performed *Bmpr2* KO rescue experiments using BMPR-2 deletion mutants.

When a gRNA-resistant full-length BMPR-2 was co-expressed, the defective dendrite remodeling by *Bmpr2* KO was fully rescued (a single primary dendrite was formed in 92% of mitral cells) **(Figure 1G)**, excluding the possibility of an off-target effect of the CRISPR/Cas9 KO. Next, we expressed a kinase domain-deleted BMPR-2 mutant (*Bmpr2ΔKinase*) combined with the *Bmpr2* KO. In this situation, mitral cells cannot mediate the canonical pathway via BMPR-2. Nevertheless, mitral cells still formed single primary dendrites (single: 95%) **(Figure 1H)**, suggesting that the canonical pathway is not needed for normal dendrite remodeling. On the other hand, a rescue experiment with a tail domain-deleted BMPR-2 mutant (*Bmpr2ΔTail*) failed to rescue the KO phenotype (single: 56%), suggesting that the non-canonical pathway through the BMPR-2 tail domain is involved in the dendrite remodeling **(Figure 1I)**.

We also examined whether ligand-binding to the BMPR-2 is required to ensure single primary dendrites. When an extracellular (EC) domain-deleted BMPR-2 mutant (*Bmpr2ΔEC*) was expressed combined with the *Bmpr2* KO, single primary dendrites were still formed (single: 83%) **(Figure 2A and B)**. This result indicates that BMPR-2 facilitates dendrite pruning without ligand-binding **(Figure 2C)**.

**Figure 2.**
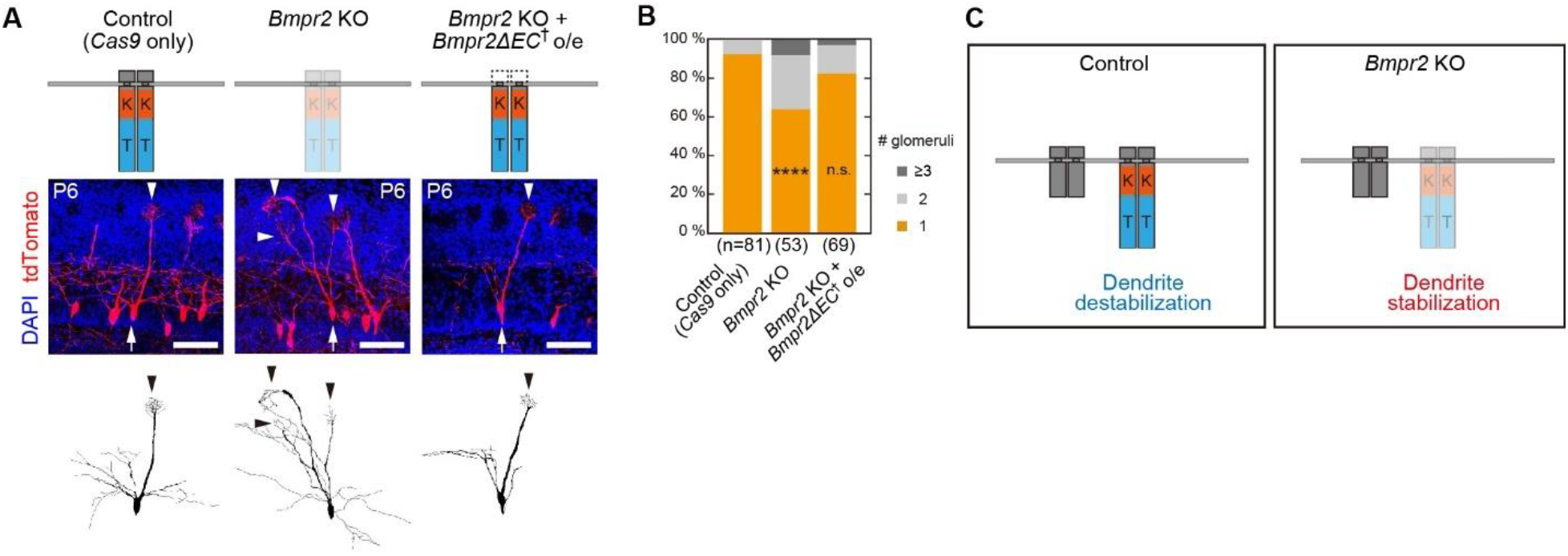
BMPR-2 facilitates dendrite destabilization without BMP ligands. **(A)** Rescue of *Bmpr2* KO using *Bmpr2ΔEC* which lacks the extracellular domain. Control and *Bmpr2* KO images are the same as in Figure 1D. Age, P6. Scale bars,100 μm. **(B)** Quantification of the number of glomeruli innervated by single mitral cells. n, number of mitral cells. n.s., non-significant. **** p<0.0001 (χ^2^ test with Bonferroni correction, compared to the control). **(C)** A schema illustrating the role of BMPR-2 in dendrite destabillization. In the *Bmpr2* KO, multiple dendrites were stabilized. However, a rescue with *Bmpr2ΔEC* showed normal dendrite destabilization, suggesting that a BMPR-2 C-terminal tail facilitates dendrite destabilization without BMP ligands.

### BMPR-2 tail domain inhibits LIMK1 to facilitate dendrite destabilization

The targets for non-canonical BMP pathways include LIM kinase (LIMK), p38/MAPK, phosphatidylinositol 3-kinase (PI3K), and Cdc42 (Gamez et al., 2013). Earlier studies have shown that BMPR-2 regulates neurite extension via LIMK (Foletta et al., 2003; Lee-Hoeflich et al., 2004; Wen et al., 2007), and the BMPR-2 tail domain was reported to capture the LIM domain of LIMK, which is required for interaction with its activators, such as Rock and PAK (Foletta et al., 2003). It is also known that active LIMK inhibits an actin depolymerization protein, cofilin, by phosphorylation, thereby stabilizing F-actin (Zebra et al., 2000). Therefore, we examined whether dendritic stability is controlled by BMPR-2-LIMK interaction during the developmental remodeling process.

We overexpressed one of the LIMKs, LIMK1, in mitral cells. We found that multiple primary dendrites are stabilized by LIMK1 overexpression, similar to the *Bmpr2* KO phenotype (single: 58%) **(Figure 3A and B)**. As LIMK is known to promote F-actin formation (Zebda et al., 2000), the formation of multiple primary dendrites is likely due to the over-stabilization of F-actin. We also found that the LIMK1 overexpression phenotype is rescued by the overexpression of BMPR-2 (single: 84%), supporting the notion that BMPR-2 inhibits LIMK1 in mitral cells **(Figure 3C)**. Rescue experiments with a series of deletion mutants (*Bmpr2ΔKinase*, *Bmpr2ΔTail*, and *Bmpr2ΔEC*) revealed that the BMPR-2 tail domain inhibits LIMK1 without the presence of BMP ligands (single: 89%, 62%, and 77%, respectively) **(Figure 3D-G)**. These results suggest that ligand-free BMPR-2 inhibits LIMK, thereby preventing the excessive stabilization of F-actin and dendrites **(Figure 3H)**.

**Figure 3.**
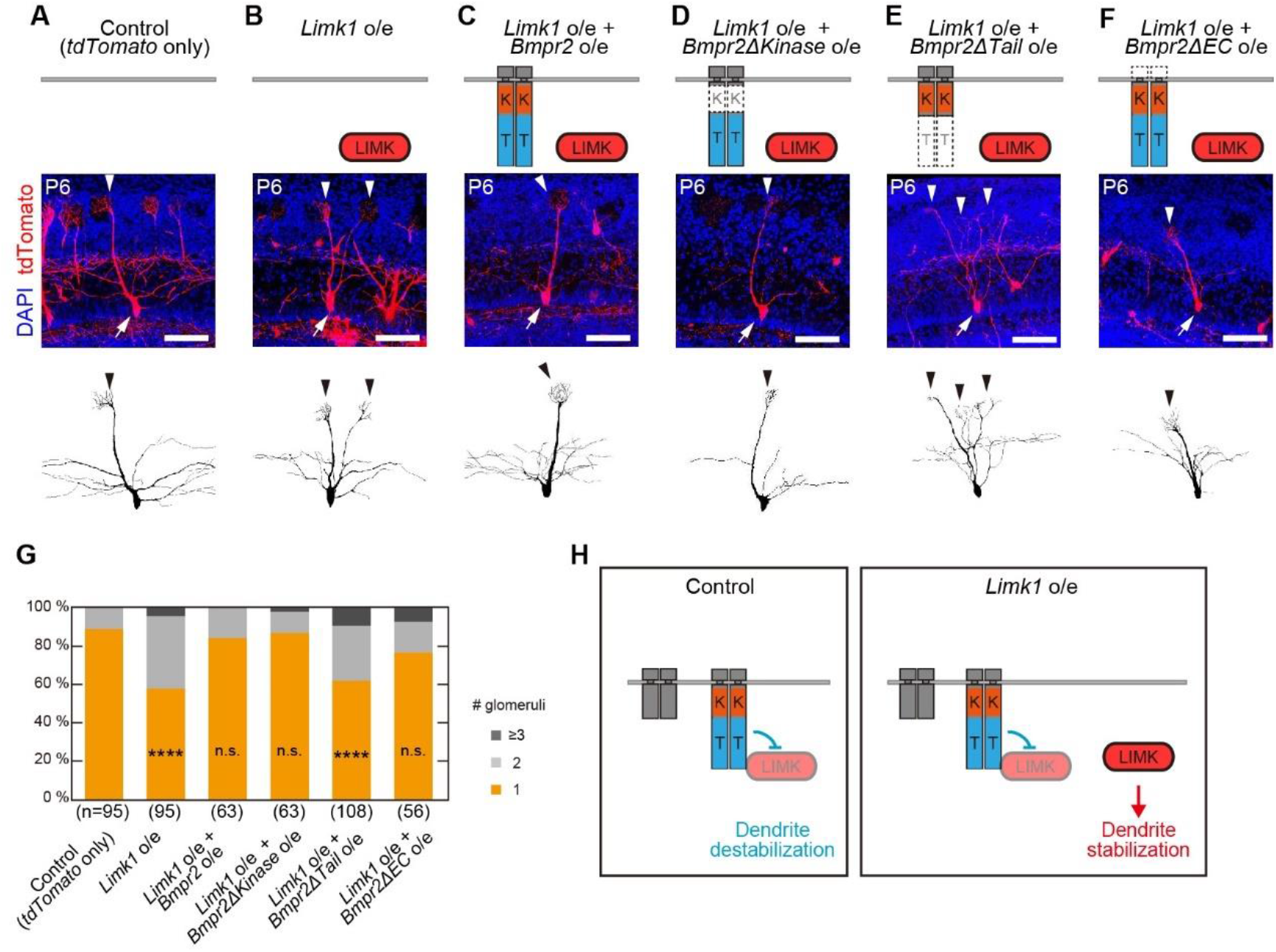
BMPR-2 tail domain inhibits LIMK1 and thereby facilitates dendrite destabilization. **(A-F)** Overexpression of *Limk1* and its rescue by *Bmpr2* deletion mutants. *Limk1* overexpression leads to the formation of multiple primary dendrites. This phenotype was suppressed by the C-terminal tail domain of BMPR-2, suggesting that the tail domain has an inhibitory role for LIMK. Age, P6. Arrows and arrowheads indicate somata and primary dendrites, respectively. Scale bars,100 μm. **(G)** Quantification of the number of glomeruli innervated per cell in (A-F). n.s., non-significant. n, number of mitral cells. **** p<0.0001 (χ^2^ test with Bonferroni correction, compared to the control). **(H)** Schematic summary of the results. *Limk1* overexpression leads to the stabilization of multiple primary dendrites in mitral cells. However, the C-terminal tail domain of BMPR-2 rescued this phenotype without ligand when overexpressed.

### BMP ligands promote dendrite stabilization

We next examined the role of BMP ligands in dendrite remodeling. In order to inhibit ligand binding to all BMPRs in mitral cells, we expressed a BMP antagonist, Noggin, in mitral cells (Groppe et al., 2002; Zimmerman et al., 1996). While the majority of mitral cells formed a single primary dendrite, 8 out of 122 mitral cells (6.6%) failed to extend any primary dendrites into the glomerular layer **(Figure 4A and B)**. We did not find this phenotype in controls. This suggests that ligand-binding to BMPRs is required to form mature primary dendrites, while BMPR-2 may not be the only BMP receptors that mediate this process.

**Figure 4.**
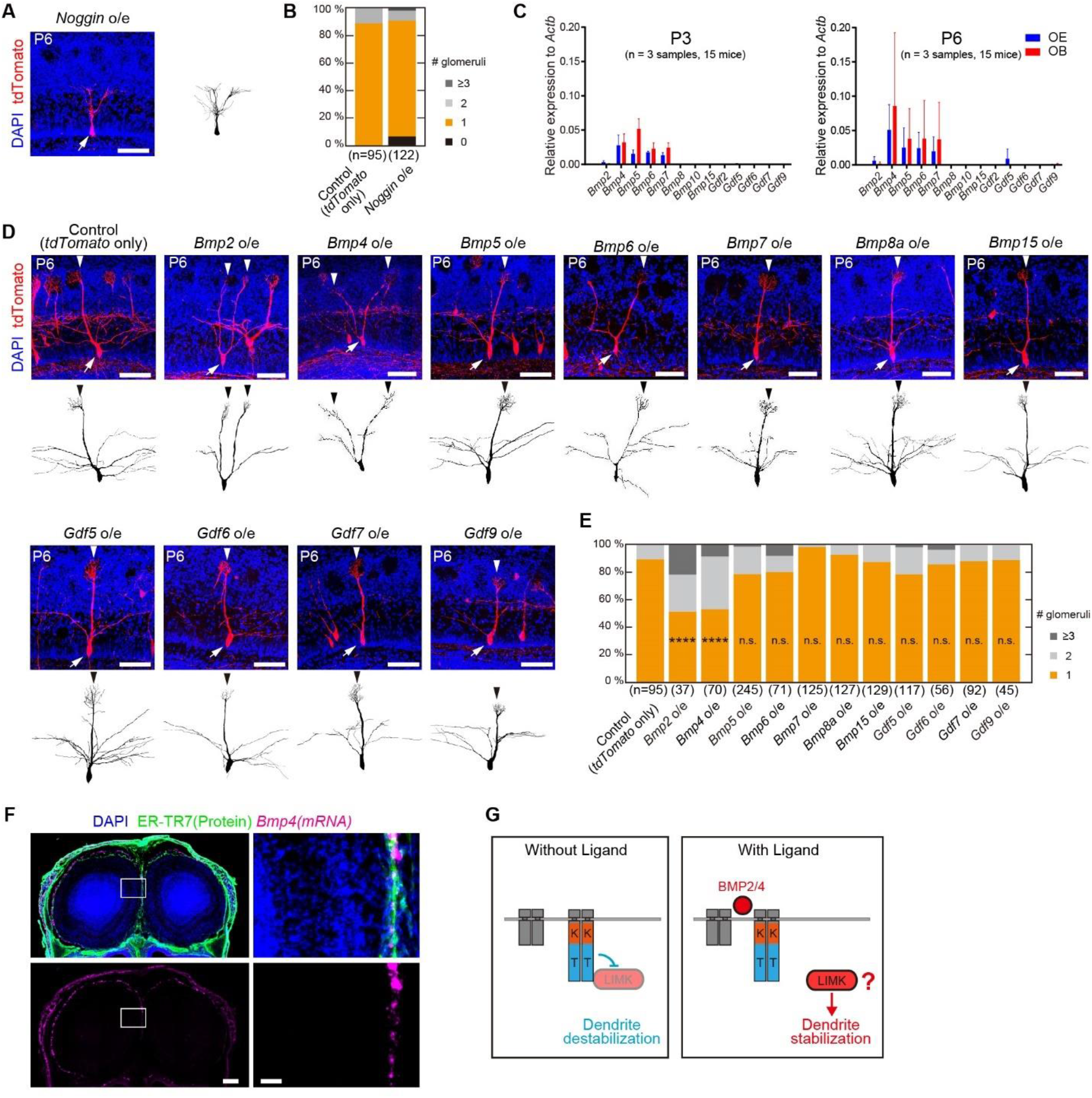
BMP2/4 stabilize dendrites. **(A)** Overexpression of the BMP antagonist, Noggin. Some mitral cells did not have any primary dendrites. Age, P6. Arrows and arrowheads indicate somata and primary dendrites, respectively. Scale bars, 100 μm. **(B)** Quantification of the number of glomeruli innervated per mitral cell. **(C)** Expression of BMP ligand genes in the OE and OB at P3 and P6. The expression levels were determined by qPCR and normalized to the *Actb* expression levels. Each cDNA pool was synthesized from mRNAs collected from 5 mice. **(D)** Overexpression of BMP and GDF genes in mitral cells. Overexpression of *Bmp2/4* led to the stabilization of multiple primary dendrites in mitral cells. The control image is the same as in Figure 3A. Age, P6. Arrows and arrowheads indicate somata and primary dendrites, respectively. Scale bars, 100 μm. **(E)** Quantification of the number of glomeruli innervated per mitral cell. n.s., non-significant. n, number of mitral cells. **** p<0.0001 (χ^2^ test with Bonferroni correction, compared to the control). **(F)** Localization of *Bmp4* mRNA was analyzed by *in situ* hybridization (RNA scope) combined with immunostaining for a meninge marker, anti-ER-TR7 immunostaining. Scale bars are 250 μm (left) and 100 μm (right). **(G)** A schematic summary of the experiment. Ligand-free BMPR-2 facilitates dendrite destabilization by inhibiting LIMK. However, when bound to BMP2/4, BMPR-2 facilitates dendrite stabilization, most likely by releasing LIMK.

BMPs are members of the transforming growth factor (TGF) –β superfamily. It has been known that BMPR-2 can potentially bind to 13 types of ligands (Mueller and Nickel, 2012). Therefore, we examined the expression of all 13 types by qPCR and found that BMP2, 4, 5, 6, 7, 8, 10, 15 and GDF2, 5, 6, 7, 9 are expressed in the olfactory epithelium (OE) and/or OB at P3 and P6 **(Figure 4C)**(Peretto et al., 2002). We then overexpressed each of these ligands in mitral cells using *in utero* electroporation, expecting autocrine and/or paracrine effects. We could not obtain BMP10 and GDF2 samples because overexpression of these ligands impaired brain development (data not shown). Among the remaining BMPs and GDFs, only BMP2 and BMP4 affected dendrite remodeling **(Figure 4D and E)**: More primary dendrites were stabilized by BMP2 and BMP4 overexpression (single: 51% and 53%, respectively). These results indicate that BMPR-2 destabilizes dendrites through LIMK inhibition without ligands, but stabilizes dendrites upon ligand binding, most likely by releasing LIMK from its tail domain **(Figure 4G)**(Foletta et al., 2003).

*In situ* hybridization of *Bmp2* and *Bmp4* mRNA revealed its localization at the surface of the OB but not the OE **(Figure S3A-D)**. Immunostaining for a meningeal marker, ER-TR7, revealed that *Bmp4* is expressed in meninges **(Figure 4F)**. *Bmp2* showed a similar pattern to *Bmp4*, but at a lower level. To examine the protein localization of BMP4, we generated *GFP::Bmp4* knock-in mice; however, we could not reliably detect the GFP signals due to technical limitations (data not shown). In general, it is extremely difficult to immunostain secreted guidance molecules. While we cannot know the localization of secreted BMP proteins, the mRNA data suggest that BMP is enriched in the surface of the OB.

### Rac1 and PAK activate LIMK1 to stabilize dendrites

How then is the free LIMK activated to stabilize dendrites? It is known that the phosphorylation of LIMK is induced by Rho-family small GTPases, such as RhoA, Cdc42 and Rac1 (Jaffe and Hall, 2005). We overexpressed these small GTPases in mitral cells and found that only Rac1 overexpression stabilizes multiple primary dendrites at P6 (single: 90%, 90%, and 45%, respectively) **(Figure 5A-D)**. Moreover, this phenotype was rescued when combined with a *Limk1* KO, suggesting that Rac1 stabilizes dendrites through LIMK1 (single: 83%) **(Figure 5E)**. We also tested PAK, an immediate downstream target of Rac1. Overexpression of *Pak1* showed a similar phenotype to the *Rac1* overexpression, showing the stabilization of multiple primary dendrites (single: 41%) **(Figure 5F)**.

**Figure 5.**
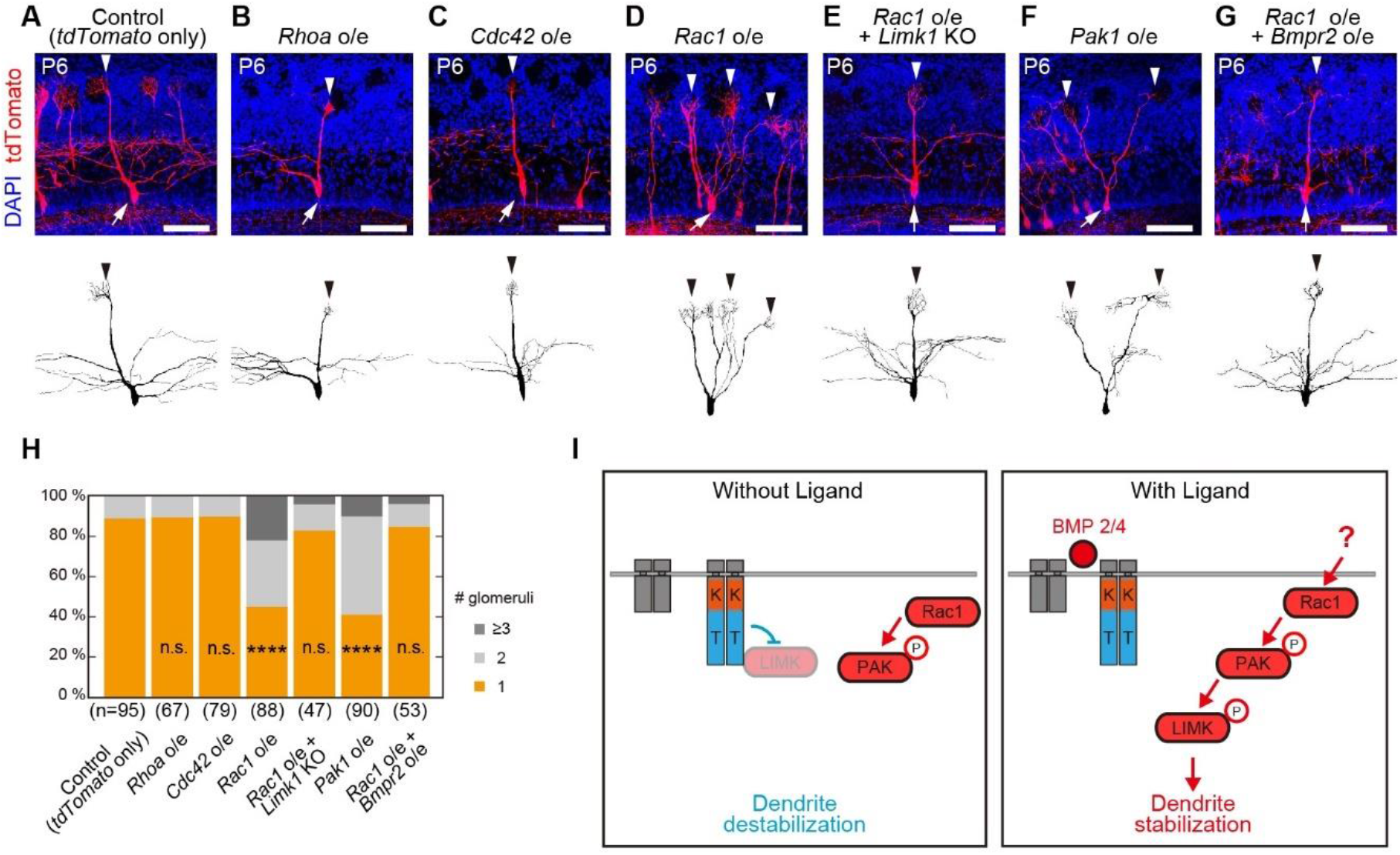
Free LIMK1 is activated by Rac1 through PAK. **(A-G)** Overexpression of the Rho-family small GTPases. Overexpression of *Rac1*, but not *Rhoa* and *Cdc42*, stabilized multiple primary dendrites. The phenotype of *Rac1* overexpression was rescued by *Limk1* KO. *Pak1* overexpression demonstrated a similar over-stabilization phenotype. The overexpression phenotype of *Rac1* was rescued by *Bmpr2* overexpression. The control image is the same as in **Figure 3A**. Age, P6. Arrows and arrowheads indicate somata and primary dendrites, respectively. Scale bars,100 μm. **(H)** Quantification of the number of glomeruli innervated per mitral cell. n.s., non-significant. **** p<0.0001 (χ^2^ test with Bonferroni correction, vs control). n, number of mitral cells. **(I)** A schematic summary of the experiments showing the activation mechanisms of LIMK. The BMP2/4-bound BMPR-2 likely releases LIMK. The free LIMK is activated by Rac1 and PAK, leading to the stabilization of primary dendrites.

Next, we examined the interplay between BMPR-2 and Rac1 in dendrite remodeling. While the overexpression of Rac1 stabilized multiple primary dendrites, the overexpression of BMPR-2 rescued this phenotype (single: 85%) **(Figure 5G)**. These results suggest that BMPR-2 has a “gating” function for Rac1 signaling: Rac1 signals are conveyed to LIMK and facilitate dendrite stabilization only when the BMP ligands bind to BMPR-2 **(Figure 5I)**.

### Glutamatergic inputs via NMDARs activates Rac1 at dendritic tufts

Rac1 is known to play a key role in long-term potentiation based on glutamatergic synaptic inputs via NMDARs in hippocampal neurons (Nakahata and Yasuda, 2018; Tu et al., 2020). We, therefore, considered the possibility that Rac1 is activated by neuronal activity in developing mitral cells. We introduced a Förster resonance energy transfer (FRET) biosensor for Rac1, RaichuEV-Rac1 (Komatsu et al., 2011), into mitral cells using *in utero* electroporation. We prepared OB slices from P3-5 mice, and FRET signals were monitored with 2-photon microscopy **(Figure 6A)**.

**Figure 6.**
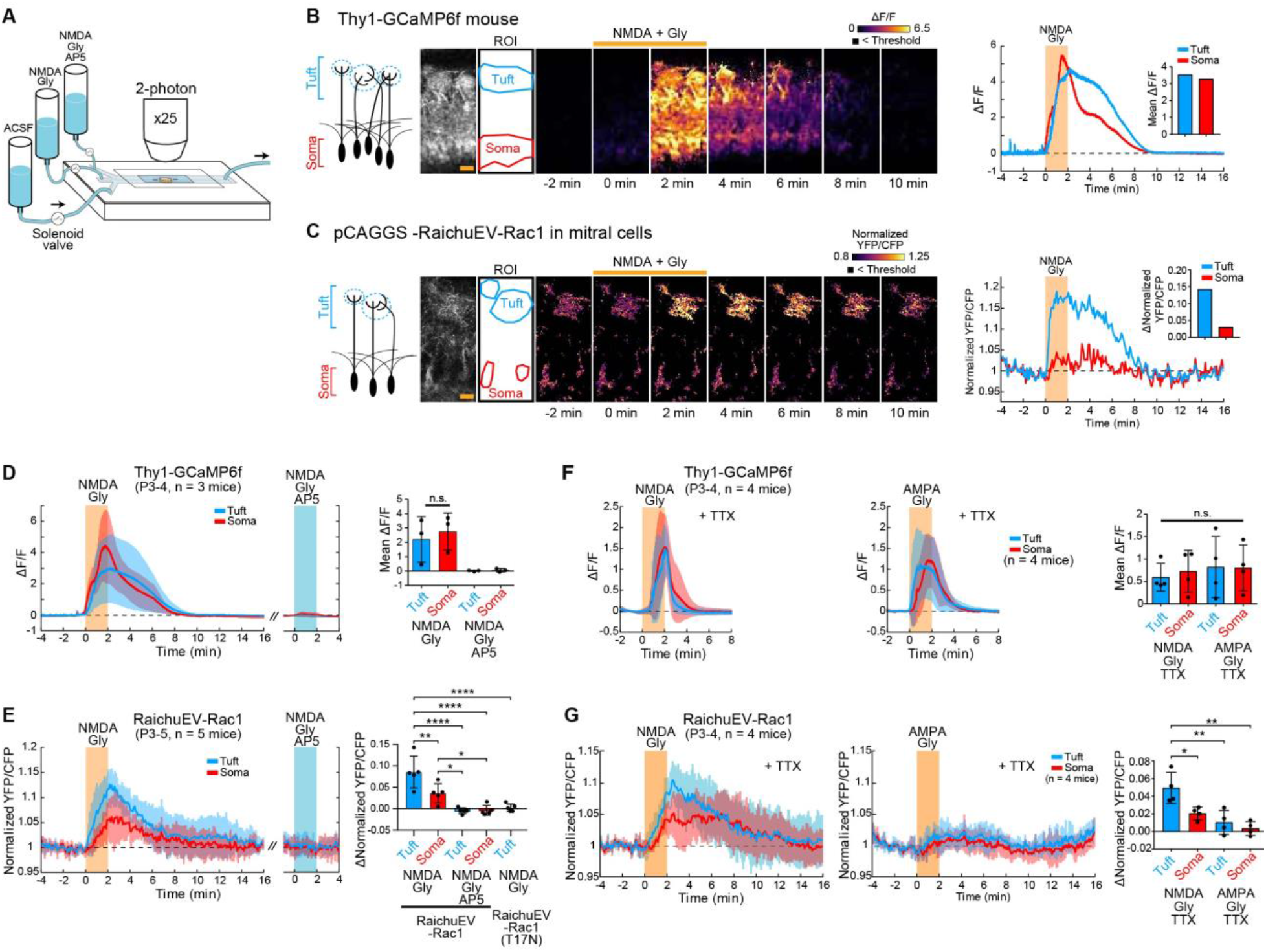
Rac1 is activated by neuronal activity via NMDAR. **(A)** Schematic diagram of the OB slice imaging. Acute OB slices were prepared from P3-5 mice. ACSF and other drug cocktails were perfused from syringes by gravity. The flow was controlled by solenoid valves. GCaMP or FRET signals were monitored by 2-photon microscopy. **(B, C)** Representative responses of Ca^2+^ and FRET signals. For Ca^2+^ imaging, Thy1-GCaMP6f transgenic mice were used, in which mitral/tufted cells are specifically labeled. For Rac1 FRET imaging, pCAGGS-RaichuEV-Rac1 was introduced into mitral/tufted cells by *in utero* electroporation. 100 μM NMDA and 40 μM Gly was applied during 0-2 min. Pixels below the threshold level of basal fluorescence are shown in black. Images were processed with a median filter. GCaMP6f signals are shown as ΔF/F values in pseudocolor. FRET signals are shown as the normalized YFP/CFP ratio. Left images show the field of view. ROIs were drawn on the tuft and soma areas, and the time courses are shown on the right. Orange shades indicate the stimulation period (2 min). Scale bars, 25 μm. **(D, E)** Average time courses of Ca^2+^ and FRET signals at tufts (blue) and somata (red) (n=3, 5 mice respectively). SD is shown by pale colors behind the traces. Orange and blue shades represent NMDA and NMDA + AP5 stimulation period, respectively. Quantification of the mean responses (0-3 min) are shown on the right. Mean ΔF/F and Δ Normalized YFP/CFP for 3 min were calculated, where the mean signals before stimulation (1 min) were used as references. An inactive mutant of RaichuEV-Rac1 (T17N) was also tested. n.s., non-significant, ** p<0.01, **** p<0.0001 (One-way ANOVA with Tukey’s post hoc test). **(F, G)** Average time courses of Ca^2+^ and FRET signals under TTX (n=4 mice each). Orange shades represent NMDA or AMPA + Gly stimulation period. Quantification of the responses are shown on the right. n.s., non-significant, * p<0.05, ** p<0.01 (One-way ANOVA with Tukey’s post hoc test).

In the control experiment, we used *Thy1-GCaMP6f* transgenic mice, in which GCaMP6f was specifically expressed in mitral/tufted cells. NMDA application (100μM) produced robust Ca^2+^ responses both in mitral cell somata and in dendritic tufts **(Figure 6B and D)**. In FRET imaging for Rac1, NMDA application also produced robust responses in mitral cells **(Figure 6C and E)**. The NMDAR antagonist, AP5 (100 μM), abolished the FRET responses to NMDA. We also confirmed that a mutant RaichuEV-Rac1 (T17N), which contains the inactive form of the Rac1 domain, did not produce any FRET responses. FRET signals were more prominent in dendritic tufts than in somata, which was not seen for GCaMP Ca^2+^ imaging **(Figure 6E)**. These results demonstrate that glutamatergic inputs through NMDARs induces Rac1 activation in developing mitral cells, especially in dendritic tufts.

We further examined how Rac1 is activated within mitral cells. We examined the responses of mitral cells to NMDA (100 μM) and AMPA (100 μM) in the presence of the sodium channel blocker, tetrodotoxin (TTX). In the control experiments with GCaMP6f, Ca^2+^ responses to NMDA and AMPA were comparable **(Figure 6F)**. In contrast, the FRET signals for Rac1 showed much stronger responses to NMDA than to AMPA, indicating that it is an NMDAR-mediated Ca^2+^ influx, rather than voltage-gated Ca^2+^ channels, that is the major source of Rac1 activation **(Figure 6G)**. This can also explain why Rac1 activation was confined to the dendritic tufts.

### F-actin formation by cofilin phosphorylation occurs preferentially at dendritic tufts

How can activated LIMK stabilize dendrites? LIMK is known to phosphorylate cofilin at S3 (Oser and Condeelis, 2009). While non-phosphorylated cofilin facilitates F-actin severing and disassembly, phosphorylated cofilin allows F-actin formation (Bernstein and Bamburg, 2010). We, therefore, tested whether cofilin controls F-actin stability and dendritic remodeling downstream of LIMK in mitral cells. When an active form cofilin mutant (phosphoblock S3A substitution, cofilin1(S3A) hereafter) was introduced by *in utero* electroporation and expressed under the CAG promoter, neuronal migration was impaired **(Figure S4A)**. We, therefore, expressed cofilin1(S3A) from P0 using the Tet-On system and doxycycline administration (P0-6). When *Limk1* alone was overexpressed, multiple primary dendrites were formed at P6; however, the expression of *cofilin1(S3A)* from P0 rescued the *Limk1* overexpression phenotype, forming single primary dendrites (single: 58% and 85%, respectively) **(Figure 7A and B)**. This result indicates that LIMK stabilizes dendrites by phosphorylating cofilin.

**Figure 7.**
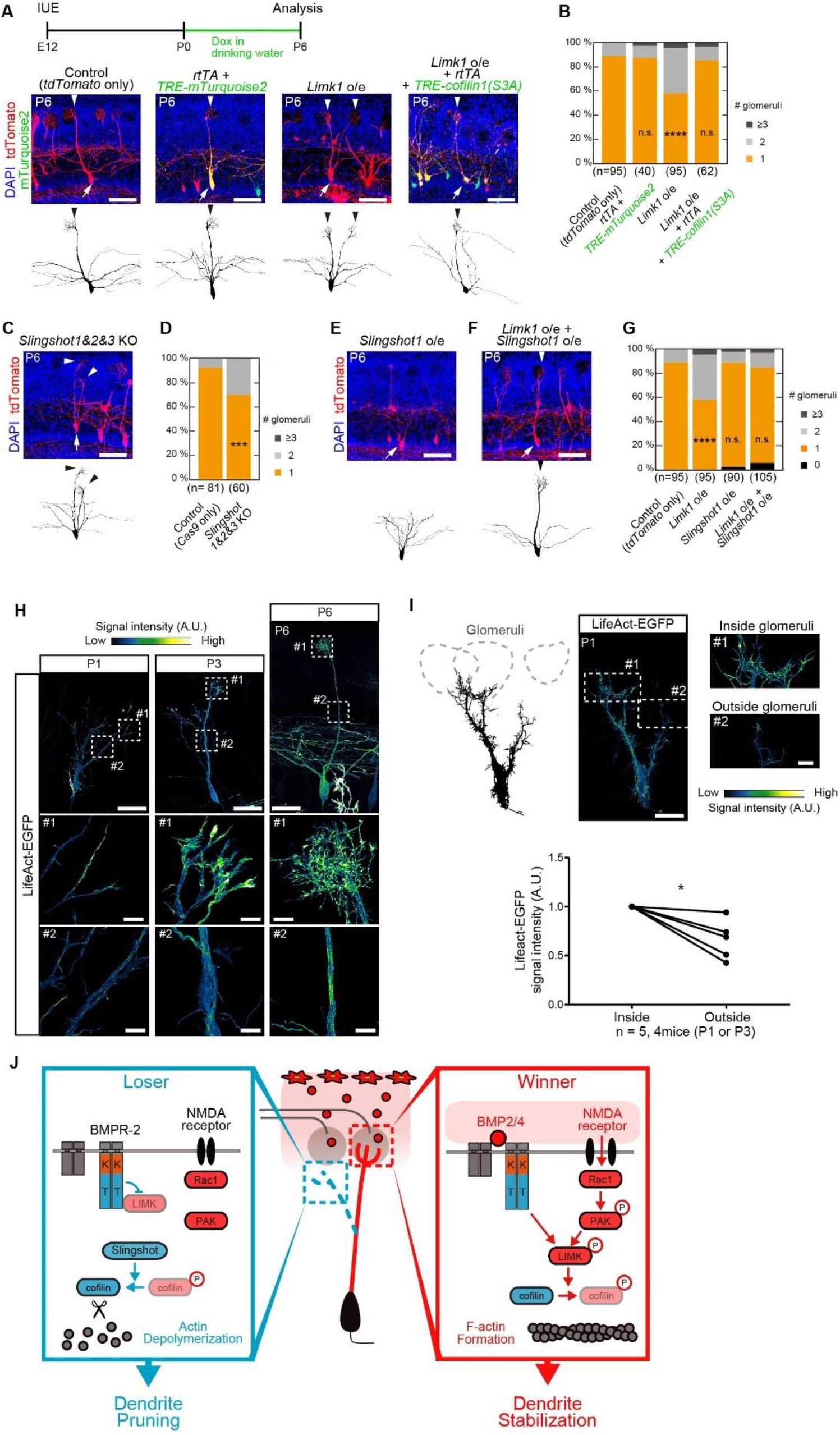
Regulation of the actin cytoskeleton by cofilin at tufts is a key for dendrite remodeling. **(A)** A phospho-blocked cofilin1 mutant, cofilin1(S3A), rescued the phenotype of *Limk1* overexpression. In this experiment, *cofilin1(S3A)* was expressed using the Tet-On system (*rtTA* and TRE-*cofilin1(S3A)*) because *cofilin1(S3A)* was found to affect neuronal migration at an earlier stage. *cofilin1(S3A)* expression was induced by doxycycline (2mg/mL) in drinking water from P0. The control and *Limk1* o/e images are the same as in **Figure 3A** and **B**. Age, P6. Arrows and arrowheads indicate somata and primary dendrites, respectively. Scale bars, 100 μm. **(B)** Quantification of the number of glomeruli innervated by single mitral cells. n, number of mitral cells. (n.s., non-significant, *** p<0.001 (χ^2^ test with Bonferroni correction, vs control). **(C)** Slingshot triple KO (*Slingshot1, 2,* and *3*) using the CRISPR/Cas9 system combined with *in utero* electroporation. The triple KO caused a pruning defect. Age, P6. Scale bars, 100 μm. **(D)** Quantification of the number of glomeruli innervated per mitral cell. n, number of mitral cells. n.s., non-significant. *** p<0.001 (χ^2^ test, vs control). **(E, F)** Overexpression of *Slingshot1*. Some mitral cells did not form any primary dendrites. The phenotype of *Limk1* overexpression was rescued by co-overexpression of *Slingshot1*. Age, P6. Arrows and arrowheads indicate somata and primary dendrites, respectively. Scale bars, 100μm. **(G)** Quantification of the number of glomeruli innervated per mitral cell. n, number of mitral cells. Mitral cells with no primary dendrites were excluded from the statistical tests. n.s., non-significant. **** p<0.0001 (χ^2^ test, with Bonferroni correction, vs control). **(H)** Visualization of F-actin with LifeAct-EGFP at different stages of dendrite remodeling. Top panels show low-magnification images. Middle and bottom images show high-magnification images of dendritic tufts and shafts, respectively. LifeAct signals are higher at dendritic tufts throughout development, while the signals gradually accumulated to shafts and somata over time. Scale bars are 50 μm (top) and 10 μm (middle and bottom). **(I)** A representative mitral cell extending dendrites to both inside and outside glomeruli (P1). Traces are shown on the left. LifeAct signals were most prominent in dendritic terminals inside glomeruli. Quantification of LifeAct-EGFP signals in primary dendrites inside and outside glomeruli are shown below. The average intensity within 30μm of the tip of the dendrites were analyzed. Additional raw images are shown in **Figure S5**. Age, P1 and P3. Scale bars are 50 μm (left) and 10 μm (right). * p<0.05 (Student’s t-test). **(J)** A proposed model of dendrite remodeling in mitral cells. Ligand-free BMPR-2 inhibits LIMK via its tail domain. In this situation cofilin is dephosphorylated by Slingshot and depolymerizes actin, leading to dendrite pruning. On the other hand, BMPR-2 releases LIMK1 upon BMP2/4 binding. The free LIMK is activated by neuronal activity via NMDAR-Rac1-PAK signaling and inhibits cofilin, leading to dendrite stabilization through F-actin formation. Thus, primary dendrites can be stabilized when BMP2/4 and glutamatergic inputs co-exist. As the BMP ligands are likely to be enriched on the OB surface, dendrites reaching the glomerular layer can be stabilized.

As LIMK stabilizes dendrites through cofilin phosphorylation, we assumed that cofilin phosphatases may play an opposing role in dendrite remodeling. Here we examined the role of Slingshot, which is known to dephosphorylate cofilin (Niwa et al., 2002). We performed a triple KO of *Slingshot* genes (*Slingshot1, 2, and 3*) using the CRISPR/Cas9 system combined with *in utero* electroporation. The *Slingshot* triple KO demonstrated impairment in dendrite pruning (single: 70%) **(Figure 7C and D)**. In contrast, *Slingshot1* overexpression caused the complete loss of primary dendrites in 2 out of 90 mitral cells **(Figure 7E and G)**, supporting the notion that Slingshot facilitates dendrite pruning. We also found that the *Limk1* overexpression phenotype is rescued by the co-expression of *Slingshot1* (single: *Limk1* 58%, *Limk1 + Slingshot1* 84%, 6/105 cells had no primary dendrite) **(Figure 7F and G)**. These results indicate that cofilin plays a pivotal role in dendrite remodeling: Phosphorylation and dephosphorylation of cofilin leads to stabilization and destabilization of primary dendrites, respectively.

Lastly, we examined whether F-actin formation and disassembly are spatially controlled during the dendrite remodeling process. To visualize the F-actin formation, we expressed an F-actin marker, LifeAct-EGFP in mitral cells. LifeAct signal was most prominent at the dendritic tips early in the development (P1-3), but it gradually spread to the dendritic shafts and somata at later stages **(Figure 7H and S4B)**. We next examined whether LifeAct signals are different in different dendrites during the remodeling process. We found that LifeAct signals are stronger at the dendritic tips extending into the glomerular layer than those retained in the external plexiform layer **(Figure 7I and S5)**. These results suggest that mitral cell dendrites receive F-actin stabilization signals, BMP ligands and glutamate, more within the glomerulus than outside.

## DISCUSSION

### Bidirectional roles of BMPR-2 in selective dendrite remodeling

Excessive neurite extension and subsequent remodeling are fundamental processes to establish mature neuronal circuits in the mammalian nervous system. However, the mechanisms underlying selective remodeling have remained largely unknown. In this study, we revealed the bidirectional role of BMPR-2 in dendrite remodeling: Ligand-free BMPR-2 facilitates dendrite destabilization, whereas ligand-bound BMPR-2 promotes dendrite stabilization. These functions are mediated by the intracellular tail domain of BMPR-2, which inhibits LIMK activity in the absence of ligand and releases it upon ligand binding. However, ligand-binding alone does not induce dendrite stabilization. The released LIMK has to be activated by neuronal activity through the NMDAR-Rac1-PAK pathway to stabilize dendrites. Activated LIMK phosphorylates cofilin and thereby facilitates F-actin formation at the dendritic tufts, thus leading to dendrite stabilization. Therefore, dendrites are stabilized only when BMP ligands and glutamatergic inputs co-exist **(Figure 7J)**. This means that BMPR-2 plays a key role in the interplay between molecular guidance and activity-dependent remodeling.

Unfortunately, we were unable to directly localize BMP ligands or activation of BMPR-2 due to technical limitations. However, since BMP2 and BMP4 are secreted from the OB surface, we assume that BMP concentrations are higher at the surface of the OB. Possibly, the extracellular matrix may help localize these proteins. On the other hand, glutamatergic inputs should occur mostly within the glomeruli. FRET imaging for Rac1 activity indicated that the NMDAR-dependent activation of Rac1 is confined to dendritic tufts within glomeruli **(Figure 6C)**. We thus propose that dendrites extending to glomeruli can be selectively stabilized, while other dendrites that do not reach the glomerular layer are destabilized. Distribution of LifeAct signals also support this idea (**Figure 7, S4. S5**). Similar kinds of interplay between molecular cues and activity must be important for various kinds of activity-dependent remodeling and plasticity in the nervous system.

### BMP receptors and regulation of actin cytoskeleton

Previous studies reported that non-canonical BMPR-2 signaling via LIMK regulates dendrite extension and arborization through ligand binding (Lee-Hoeflich et al., 2004; Saxena et al., 2018). However, our current study demonstrated that BMPR-2, in fact, has bidirectional roles in dendrite remodeling: The ligand-free BMPR-2 induces dendrite destabilization and the ligand-bound BMPR-2 facilitates dendrite stabilization. Thus, BMPR-2 functions as a gatekeeper for dendrite stabilization signals triggered by neuronal activity in a ligand-dependent manner.

There have been conflicting results regarding the roles of BMPR-2-LIMK1 interaction. One study showed that BMPR-2 inhibits LIMK1 activity, whereas another study demonstrated that BMPR-2 facilitates LIMK1 activation via Cdc42 (Foletta et al., 2003; Lee-Hoeflich et al., 2004). They performed similar *in vitro* kinase assays for LIMK1 using cofilin as a substrate, but the former used a full-length BMPR-2, whereas the latter used a part of the BMPR-2 tail domain. We showed that full-length BMPR-2 can rescue the phenotype of LIMK1 overexpression, supporting the former study. It is possible that the latter study used a truncated tail domain, which, in fact, has lost the ability to inhibit LIMK. Also, the LIMK binding region (751-813aa) identified in the latter study seems to play little role in our system *in vivo*: The C-terminal tail domain, rather than 751-813aa, is critical for LIMK regulation **(Figure S2B and C)**.

The regulation of actin cytoskeleton has been extensively studied in growth cone extension and dendritic spine enlargement (Bosch and Hayashi, 2012; Luo, 2002). It has also been reported that F-actin regulation underlies dendrite development such as extension and branching (Nithianandam and Chien, 2018; Tasaka et al., 2012). However, it has not been fully understood how the actin cytoskeleton is regulated during the dendrite remodeling process. In this study, we showed that the overexpression of LIMK1 causes dendrite stabilization by phosphorylating cofilin. This indicates that F-actin formation leads to dendrite stabilization. On the other hand, the severing and depolymerization of F-actin induced by a cofilin phosphatase Slingshot seems to facilitate dendrite pruning. Together, our results indicate that the regulation of the actin cytoskeleton plays a key role in dendrite remodeling. This is in stark contrast to the axon pruning process known to play a role in the remodeling of neuromuscular junction, in which the destabilization of microtubules plays a key role (Brill et al., 2016). Our finding is also different from dendrite pruning during *Drosophila* metamorphosis, in which local endocytosis at the dendritic neck plays a key role (Kanamori et al., 2015).

### Inter-neuronal and intra-neuronal competition in circuit remodeling

In the neuromuscular junction and cerebellar climbing fiber – Purkinje cell synapse, multiple axons compete toward one target. In other words, there is a competition among different axons from different neurons. Recent studies have identified positive and negative regulators of the remodeling of climbing fiber – Purkinje cell synapses (Choo et al., 2017; Hashimoto et al., 2011; Kakegawa et al., 2015; Mikuni et al., 2013; Uesaka et al., 2018; Uesaka et al., 2014). However, it remains unclear how some axons are selectively stabilized, and others are eliminated. It also remains unclear how only single winner is maintained.

In the olfactory bulb, each glomerulus is innervated by 20-50 mitral/tufted cells; however, each of mitral/tufted cells connects just one primary dendrite to a glomerulus. This indicates that there is an intra-neuronal competition, in which different dendrites from the same neuron compete to each other to become a winner. In previous studies, positive and negative regulators for dendrite morphogenesis in mitral cells have been identified (Imamura and Greer, 2009; Inoue et al., 2018; Muroyama et al., 2016); however, they cannot fully explain the mechanisms of selective remodeling. In this study, we show how some dendrites are selectively stabilized when BMP ligands and neuronal activity co-exist.

Currently, our results still cannot explain how other dendrites are pruned during the remodeling process. Ligand-free BMPR-2 only has a permissive role for dendrite pruning. There must be opposing signals to prune supernumerary dendrites (Fujimoto et al., 2019). Slingshot is one of these molecules; however, it remains unclear how they are regulated (Ohashi, 2015). In the future, it will be important to study how pruning is controlled and what mediates the competition during the neurite remodeling process.

## ACKNOWLEDGMENTS

We thank K. Svoboda (*Thy1-GCaMP6f*) for mice; P. Soriano (*pPGKFLPobpA*), F. Matsuzaki (*pCAX-Cas9* and gRNA backbone vector), and K. Aoki (*pCAGGS-RaichuEV-Rac1*) for plasmids; Y. Tsunekawa, F. Matsuzaki, H. Inomata, S. Okabe, N. Nakagawa, H. Murakoshi, and R. Yasuda for valuable technical advice; M. Nishihara, T. Ohmine, and The Research Support Center, Research Center for Human Disease Modeling, Kyushu University Graduate School of Medical Sciences, for technical assistance; M.N. Leiwe for comments on the manuscript.

Imaging experiments were in part supported by the RIKEN Kobe Light Microscopy Facility and animal experiments were in part supported by the Laboratory for Animal Resources and Genetic Engineering at the RIKEN Center for Life Science Technologies. This work was supported by grants from the PRESTO program of the Japan Science and Technology Agency (JST) (T.I.), the JSPS KAKENHI (23680038, 16H06456, and 17H06261 to T.I., 15K14327, 17K14944, and 19K06886 to S.F., 18J10215 to S.A.), The Mochida Memorial Foundation for Medical and Pharmaceutical Research, and RIKEN CDB intramural grant (T.I.). S.A. was a Junior Research Associate at RIKEN and a predoctoral research fellow of JSPS.

## AUTHOR CONTRIBUTIONS

S.A. performed most of the experiments. S.A and R.S. optimized CRISPR/Cas9-based KO system combined with *in utero* electroporation. S.F. performed microarray experiments and helped *in utero* electroporation. S.A. and S.F. performed Ca^2+^ imaging and FRET imaging of OB slices. T.I. supervised the project. S.A. and T.I. wrote the manuscript.

## METHODS

### Mice

All animal experiments were approved by the Institutional Animal Care and Use Committee (IACUC) of the RIKEN Kobe Branch and Kyushu University. *Thy1-GCaMP6f* Tg (line GP5.11) (JAX #024339) (Dana et al., 2014) has been described previously. ICR mice (Japan SLC, RRID: MGI: 5652524) were used for *in utero* electroporation. For anatomical and histological analyses, mice were deeply anesthetized by intraperitoneal injection of overdose Nembutal (Dainippon Sumitomo Pharma) or Somnopentyl (Kyoritsu Seiyaku) and perfused with phosphate buffered saline (PBS) for *in utero* electroporation samples and 4% paraformaldehyde (PFA) in PBS for other samples. Dissected brains were post-fixed with 4% PFA in PBS at 4°C overnight. For inducible gene expression by the Tet-On system, drinking water containing doxycycline (2mg/mL, Sigma-Aldrich, # D9891) and sucrose (10 % w/v) was administrated from P0.

### CRISPR/Cas9

A procedure for gRNA construction is shown in **Figure S1A**. pCAX-Cas9 and gRNA backbone vector were kind gifts from F. Matsuzaki (Tsunekawa et al., 2016). For the CRISPR/Cas9-based KO system, the three gRNAs that target different exons were designed to increase KO efficiency (Sunagawa et al., 2016). The gRNA sequences were designed by Optimized gRNA design (https://zlab.bio/guide-design-resources) or CHOPCHOP (https://chopchop.cbu.uib.no/). The gRNA sequences are shown in **Table S1**. Target sequences were amplified with forward and reverse oligonucleotides by PCR and inserted into the gRNA backbone vector at AflII sites (Tsunekawa et al., 2016).

### Plasmids

ORF sequences were amplified by PCR from P6 mouse OB or OE cDNA. Because *Gdf7* contains GC rich sequences, we could not amplify the ORF from cDNA. Instead, we designed a synthetic *Gdf7* gene with lower GC contents (GeneArt, ThermoFisher). Amplified genes were subcloned into *pCAG, pCA-FNF, or pTRE* vector. FLPo (addgene #13793, a gift from P. Soriano) was amplified by PCR and subcloned into a pCAG vector. Truncated versions of *Bmpr2* (NM_007561.4) encode the following amino acid sequences; *Bmpr2ΔKinase* (1-201 and 501-1038 aa.), *Bmpr2ΔTail* (1-529 aa.), *Bmpr2ΔEC* (1-26 and 151-1038 aa.), and *Bmpr2Δ751-813aa* (1-750 and 814-1038 aa.). gRNA-resistant *Bmpr2* (Designated *Bmpr2^†^*) was generated by PCR-mediated mutagenesis and has the following sequences on the gRNA-targeting sequences; CaagtCTaCAcaGaCCaTTcaGa, agtAAgGGatcaACaTGcTAcGG, and aGgTAtGGtGCtGTtTAcAAgGG (small characters are substituted nucleotides). A FRET sensor, RaichuEV-Rac1 (Komatsu et al., 2011), is a kind gift from K. Aoki. Newly-generated plasmids will be deposited to Addgene.

### *In utero* electroporation

To sparsely label neurons, pCAG-Flpo (Addgene #125576, 3-10 ng/μL) and pCAFNF-tdTomato (Addgene#125575, 1 μg/μL) were used. For CRISPR/Cas9-based KO, three types of gRNA plasmids (0.1 μg/μL each) and pCAX-Cas9 (0.1 μg/targeted gene/μL) were used (Tsunekawa et al., 2016). For other plasmids, concentration was 1 μg/μL (see **Table S4** for detailed conditions). *In utero* electroporation was performed as described previously (Fujimoto et al., 2019; Muroyama et al., 2016). Pregnant mice carrying E12 embryos were anesthetized with ketamine (64-80 mg/kg) and xylazine (11.2-14 mg/kg). The uterine horns were exposed by abdominal incision. A plasmid cocktail was injected into the lateral ventricle of the embryos using a glass capillary. Electric pulses (a 10ms single poration pulse at 72V followed by five 50 ms duration, 34V driving pulses with a 950 ms interval) were delivered by a CUY21EX electroporator (BEX, # CUY21EX) and forcep-type electrodes (3 mm diameter, #LF650P3, BEX). After the surgery, anesthetized mice were kept on the heating pad (IKEDA scientific, #IP-4530) until they wake up.

### Microarray

Mitral cells were labeled with EYFP using *in utero* electroporation. At P3 or P6, the OB was minced in Ca^2+^-free Ringer’s solution (138mM NaCl, 2mM MgCl_2_, 2mM sodium pyruvate, 9.4mM glucose, 2mM EGTA and 5mM HEPES, pH7.4). Minced OB was incubated with 0.88U/mL Dispase (Invitrogen), 200 U/mL Collagenase type II, and 0.1 mg/mL DNase I (Roche) for 20 minutes at 37 ℃. The sample was centrifuged at 1k rpm for 5 minutes and the pellet was resuspended in Ca^2+^ free Ringer’s solution. After repeating this washing procedure again, 0.1% bovine serum albumin and 0.05 mg/mL DNase solution were mixed gently. The sample was placed on a coverslip and the EYFP-expressing mitral cells were collected manually with a glass capillary, a micro-manipulator, TransferMan NK2 (Eppendolf), and a microinjector, CellTram vario (Eppendolf), equipped to an inverted fluorescence microscope, DMI6000B (Leica). The picked single cell was rinsed in a drop of 0.1% bovine serum albumin/ Ringer’s solution (Ca^2+^ free Ringer’s solution contained 2mM CaCl_2_), transferred to 0.4 μL of RNase free H_2_O in PCR tube, snap-frozen in liquid nitrogen, and stored at −80 ℃ until use. Single-cell cDNA synthesis and amplification was performed as described previously (Imai et al., 2009). Briefly, a T7 promoter was attached to the 3’-end and the total cDNA was PCR-amplified (30 cycles). The quality of PCR-amplified cDNA was assessed by agarose gel electrophoresis. Samples with sufficient yields, containing *EYFP* and *pcdh21* transcripts (determined by secondary PCR for *EYFP* and *pcdh21*) were used in subsequent analyses. Samples from 10 neurons were pooled for each hybridization experiment. Cy3 labeling was performed with T7 RNA polymerase using Agilent Low RNA Input Linear Amplification Kit following manufacturer’s instructions (Agilent). The microarray was performed using the SuperPrint G3 Mouse GE 8×60K Microarray Kit (Agilent, #G4852A) and a DNA microarray scanner, G2505C (Agilent). The data were analyzed with Microsoft Excel. Raw data are being submitted to the NCBI Gene Expression Omnibus (GEO) with accession number, #XXX.

### Clearing with SeeDB2

PFA-fixed brains were rinsed in PBS and then embedded in 2 % agarose gel. Samples were then sliced by a microslicer (Dosaka EM, # PRO7) at 2 mm thickness for CRISPR/Cas9 screening, 0.5 mm thickness for LifeAct samples, and 1 mm thickness for the others. Brain slices were then stained with DAPI and cleared with SeeDB2G as described previously (Ke and Imai, 2018; Ke et al., 2016). For immunostaining, slices were incubated in blocking solution (2% saponin, 0.25% fish gelatin, 0.5% skim milk, 0.5% TritonX-100 and 0.05% sodium azide in PBS) overnight. Slices were then incubated with primary antibody diluted in the blocking solution for 2 days at room temperature with gentle rotation. Anti-Tbr2 (abcam, #ab23345, 1:250) and Tbx21 (abcam, #ab91109, 1:250) were used. For Tbx21 staining, brains were treated by autoclave at 105℃ for 1 min before slicing, and rabbit anti-GFP (Invitrogen, #A11122, 1:250) was also used to label GFP positive neurons. Slices were then washed with 0.1% TritonX-100 / PBS for 1.5 hours 3 times and incubated with Alexa Fluor 555-conjugated donkey anti-rabbit IgG (1:250, ThermoFisher, #A-31572) and/or Alexa Fluor 555-conjugated donkey anti-mouse IgG (1:250, ThermoFisher, #A-31570) diluted in the blocking solution overnight with gentle rotation. After washing twice with 0.1% TritonX-100 / PBS for 2 hours, slices were incubated in Omnipaque 350 (Daiichi-Sankyo) for clearing overnight with gentle rotation.

### Confocal imaging and image processing

Cleared samples were mounted on glass slides with 0.5, 1, or 2 mm thick silicone rubber spacer as described (Ke and Imai, 2018). Samples were imaged with a 2-photon microscope (Olympus, FV1000MPE) for initial KO screening, and an inverted confocal microscope (Leica, SP8) for others. A water-immersion 25x objective lens (Olympus, XLPLN25XWMP, NA 1.05) was used for 2-photon imaging. A multi-immersion 20x (HC PL APO 20x/0.75 IMM CORR CS2, NA 0.75) and a glycerin-immersion 63x (HC PL APO 63x/1.3 Gly CORR CS2, NA 1.3) objective lenses were used for confocal imaging. High resolution images of LifeAct signals were acquired with HyVolution package of a confocal microscope Leica TCS SP8 (Leica Microsystems) and Huygens software (Scientific Volume Imaging).

### Quantification of neuronal morphology

Neuronal morphology was analyzed with Neurolucida software (MBF Bioscience). Neurons labeled with tdTomato in the mitral cell layer were analyzed. Analysis was confined to mitral cells located in the medial side of the OB. We sometimes found neurons with small soma and neurites with dendritic spines that did not contact a glomerulus; these are granule cells and were excluded from subsequent analysis. Glomeruli were identified with DAPI staining. Neuronal tracing and analysis were not blinded; however, we comprehensively analyzed all the neurons in a volume that meets the above criteria to avoid any biases in quantification.

### qPCR

Total RNA was extracted from OE and OB samples at P3 and P6 using the RNeasy Plus Mini Kit (Qiagen, #74134). In order to collect sufficient amounts of RNA, samples from 5 mice were pooled for RNA extraction. In total, 3 pooled RNA samples from 15 mice were prepared. cDNAs were synthesized with SuperScript IV (ThermoFisher, #18090010) using Oligo(dT)20 primers (ThermoFisher, #18418020). qPCR primers are described in **Table S2**. For real-time PCR, PowerUP SYBR Green (ThermoFisher, #A25742) and ABI7500 (Applied Biosystems) was used. PCR efficiencies of each primer were estimated by qPCR of plasmids encoding each target gene. Data were analyzed using Excel (Microsoft). We calculated amounts of mRNA based on Ct value and PCR efficiency. The calculated values were normalized by *Actb* gene.

### *In situ* hybridization (RNA scope)

RNA scope was used for *in situ* hybridization following manufacturer’s instructions. *Bmp2* (#406661) and *Bmp4* (#401301) probes were purchased from Advanced Cell Diagnostics. To avoid tissue shrinkage during cryoprotection, 15% fructose was used instead of 30% sucrose. In double staining experiments, conventional immunohistochemistry was performed after RNA scope. Briefly, after Amp6 of RNA scope 2.5HD Reagent kit Brown (Advanced Cell Diagnostics, #322300), sections were incubated with AlexaFluor555-Tyramide reagent (Thermo Fisher Scientific, #B40923). After washing in PBS, sections were fixed with 4% PFA/PBS for 15 min, washed in PBS, and blocked with 5% donkey serum for 30 min. Then sections were incubated with anti-ER-TR7 (1:200, abcam, # ab51824) for 1 hour at room temperature and then washed three times in PBS. Finally, sections were incubated with DAPI (1:200, DOJINDO, # 340-07971) and a secondary antibody conjugated with AlexaFruol 488 (1:200, Thermo Fisher Scientific, # A-21208) for 1 hour at room temperature. After washing, sections were mounted using ProLong Gold (Thermo Fisher Scientific, #P36930). Leica DMI6000B was used for image acquisition.

### Ca^2+^ and FRET imaging

Thy1-GCaMP6f mice were used for Ca^2+^ imaging. For FRET imaging, pCAG-RaichuEV-Rac1 (Komatsu et al., 2011) was introduced at E12 by *in utero* electroporation. FRET and Ca^2+^ imaging was performed at P3-5. Mice were anesthetized on ice and decapitated. The brain was immediately harvested and placed in cold artificial corticospinal fluid (ACSF: 125 mM NaCl, 3 mM KCl, 1.25 mM NaH_2_PO_4_, 2 mM CaCl_2_, 1 mM MgCl_2_, 25 mM NaHCO_3_, and 25 mM glucose). The brain was embedded in 2% agarose gel and sliced using a microslicer at 300 μm thickness. An OB slice was placed on a custom-made silicone chamber (Fujimoto et al., 2019) and fixed with a nylon mesh (Warner Instrument, # 64-0198). The chamber was set under a 2-photon microscope (Olympus, FV1000MPE) equipped with a water-immersion 25x objective lens (Olympus, XLPLN25XWMP, NA = 1.05, WD = 2.0). Before the imaging, the OB slice was perfused with oxygenized ACSF at least for 2 hours at 27 ℃ for recovery. The excitation laser was tuned to 920 nm for Ca^2+^ imaging and 840 nm for FRET imaging. Dichroic mirrors FV10-MRVGR/XR (Olympus, #FP1NDF4VGRXR) and FV10-MRC/YW (Olympus, #FP1NDF4CY-G) were used for Ca^2+^ and FRET imaging, respectively. Emitted signals were detected with GaAsP detectors. Images were acquired every 0.5 or 5 seconds for Ca^2+^ or FRET imaging respectively. The resolution of Ca^2+^ and FRET imaging was 1.988 and 0.497 μm/pixel respectively. Following drugs were used: 100 μM NMDA (Nacalai, Cat #22034-1), 100 μM AP5 (Sigma-Aldrich, #A5282), 100 μM AMPA (TOCRIS, #0169), 1 μM TTX (abcam, #ab120055), and 40 μM Glycine (Sigma, #G7126-100G). Glycine was co-applied with NMDA and AMPA. TTX was applied 7-20 min before NMDA or AMPA stimulation. Samples drifted by the drug application were excluded from subsequent data analysis. Data were analyzed with ImageJ. Briefly, small drifts by drug application were corrected by ImageJ plugin (Correct 3D drift) when possible. Background signals were removed by thresholding. For the analysis of Ca^2+^ imaging data, the F0 was calculated as the average before stimulation (2 or 4 min). For the FRET data, the YFP/CFP ratio was calculated at each frame and then normalized by the average value before stimulation at each pixel to create representative images, or, at the ROI level for time courses and quantification. To calculate the averaged ΔF/F and Δ normalized YFP/CFP, a mean value for 3 minutes after stimulation onset was subtracted by the mean value for 1 minute before stimulation onset.

### Statistical analysis

Excel and Prism7 were used for statistical analysis. Sample sizes for dendrite quantification were determined based on pilot experiments (using G Power). The number of neurons is described within figures, and number of animals is described in **Table S3**. χ^2^-tests were used in **Figure 1–5, 7, and S2**. Mitral cells with no primary dendrites were excluded from the statistical tests because χ^2^-tests compared the single vs. multiple dendrite population. For the multiple comparison of χ^2^-tests, Bonferroni correction was used. A One-way ANOVA was used in **Figure 6G, I, K, and N** with Tukey’s post-hoc test for multiple comparison. Student’s t-test was used in **Figure 7I**, **S1C** and **E**. Data inclusion/exclusion criteria are described in each method section.

## DATA AND SOFTWARE AVAILABILITY

All the image data will be deposited to SSBD:repository (http://ssbd.qbic.riken.jp/repository/). Microarray data will be deposited to NCBI GEO. Neurolucida tracing data will be deposited to NeuroMorpho.Org (http://neuromorpho.org/). Numerical data for all graphs are included in **Table S3**. No new program codes were generated in this study. Requests for additional data should be directed to and will be fulfilled on reasonable request by the Lead Contact, Takeshi Imai.

## SUPPLEMENTAL FIGURES

**Figure S1.**
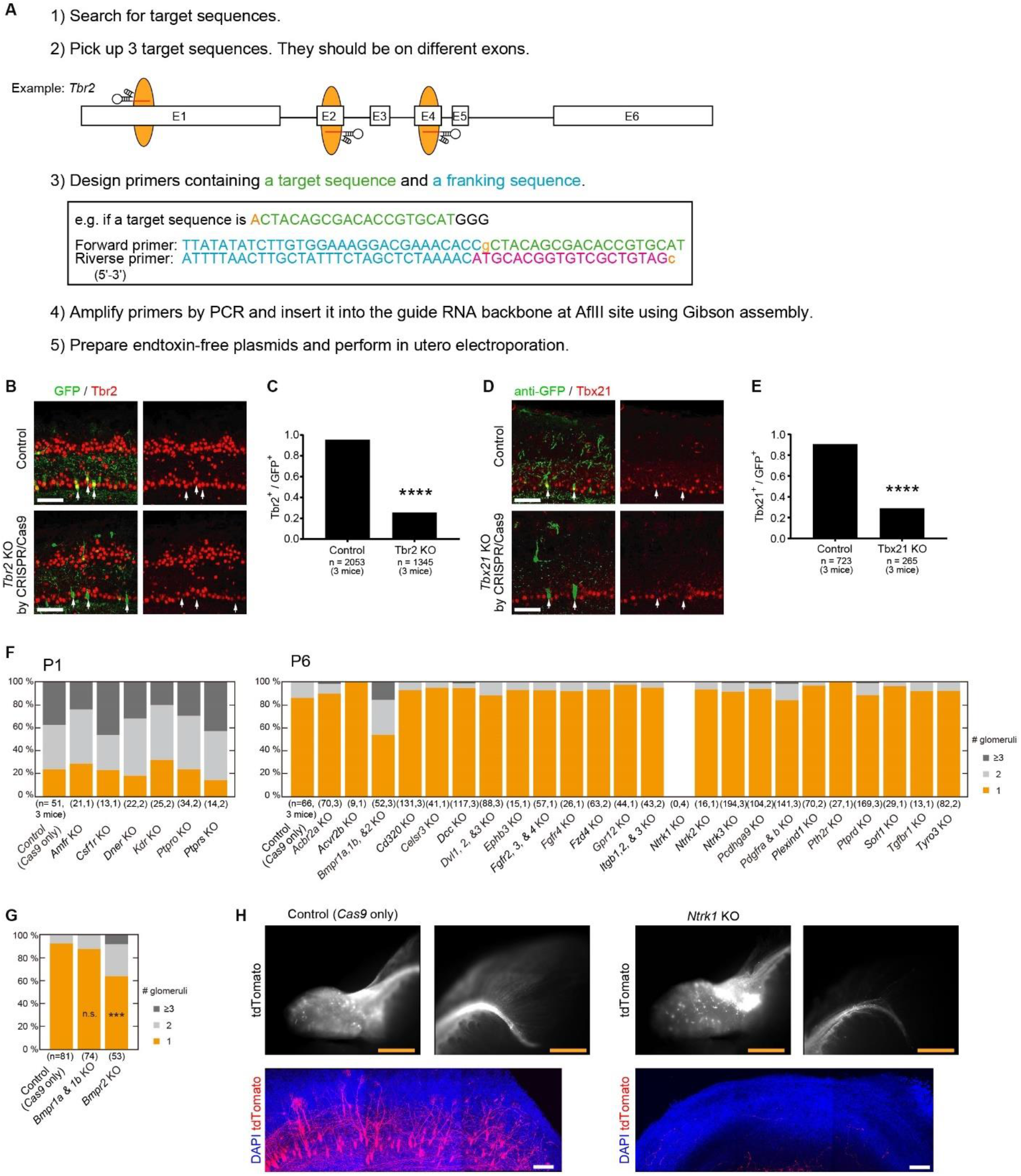
CRISPR/Cas9-based knock-out screening. **(A)** A procedure for gRNA construction. It takes approximately 4 days to prepare a gRNA plasmid. **(B-E)** The efficiency of our CRISPR/Cas9 KO screening. We have chosen Tbr2 and Tbx21, because they are specifically expressed in mitral/tufted cells and their antibodies are available. For both genes, we have designed gRNAs in the coding region as described in **(A)**. We have automatically picked the 3 best gRNAs as suggested by the gRNA designing program. An EGFP expression vector was co-expressed with CRISPR/Cas9 plasmids. Immunostaining demonstrated a KO efficiency of 70.6% on average (73% and 68%, respectively). **** p<0.001 (χ^2^ test). Scale bars, 100 μm. **(F)** Candidate cell surface receptors tested in this study. Based on the single-cell microarray data for mitral cells, we have chosen cell surface receptors. We performed CRISPR/Cas9-based KO screening for these receptor genes in single, double, or triple KOs in the initial screening. We could not find mitral cell somata for *Ntkr1* KO samples, while labeled axons were found. It is possible that retrograde NGF signals received by Trk-A (encoded by *Ntkr1*) are essential for mitral cell survival (Ginty and Segal, 2002). **(G)** After identifying defective dendrite pruning in the *Bmpr1a/1b/2* triple KO experiment, we performed further KO experiments, and identified *Bmpr2* as a critical gene for dendrite remodeling. n.s., non-significant, *** p<0.001 (χ2 test with Bonferroni correction, vs control). **(H)** *Ntrk1* KO. Top panels show epifluorescence images of OB and mitral cell axons in the lateral olfactory tract. Bottom panels are 400 μm z-stack OB images acquired with 2-photon microscopy. Top and bottom are from the same samples. Axons were observed, but mitral cell somata were not seen in the OB. Scale bars, 1 mm (Top) and 100 μm (bottom).

**Figure S2.**
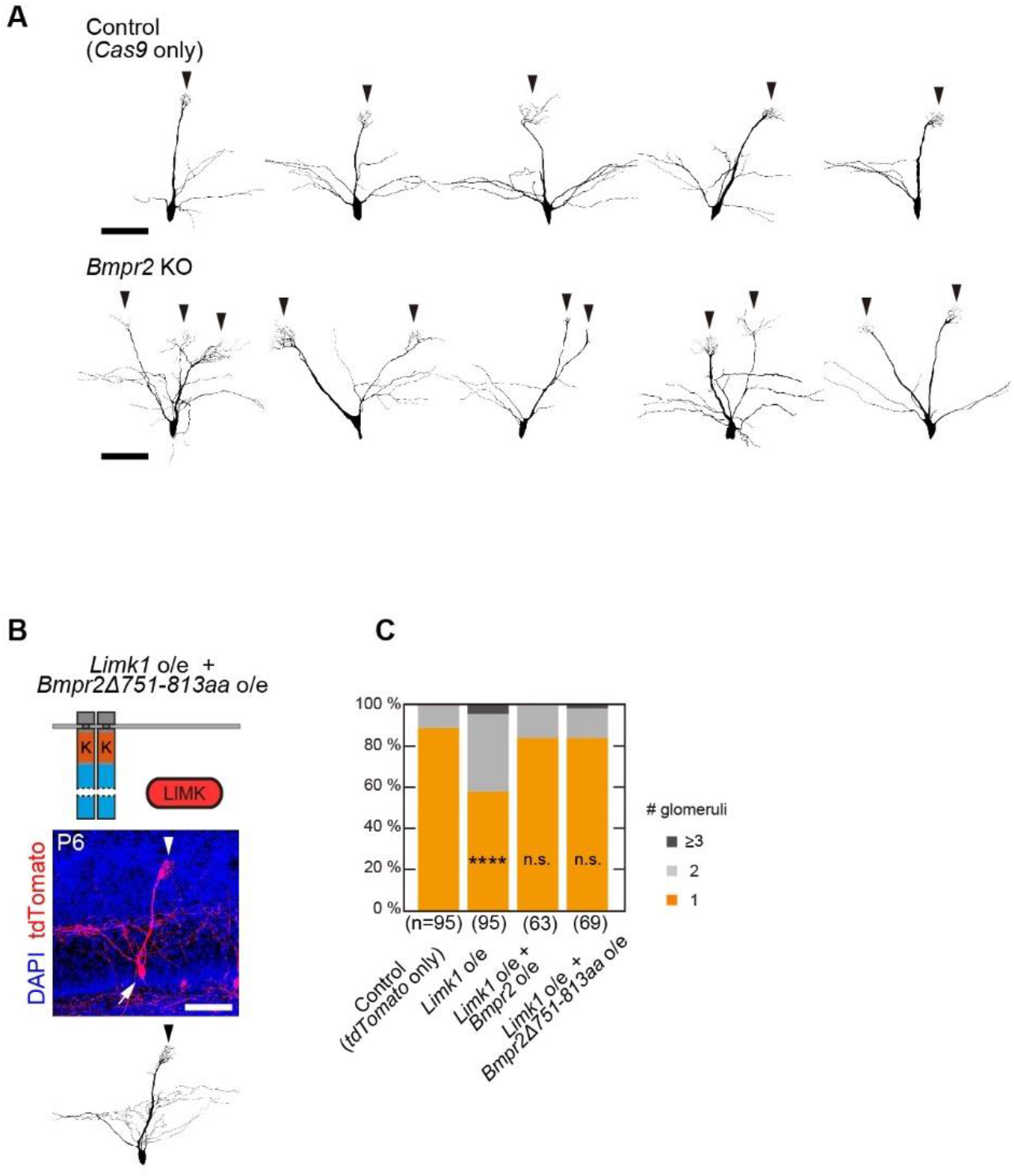
A rescue experiment of *Limk1* overexpression with *Bmpr2Δ751-813aa*. **(A)** Representative traces of control and *Bmpr2* KO mitral cells at P6. Scale bars, 100 μm. **(B)** A previous study reported the presence of “LIMK-binding regions” (751-813aa) (Lee-Hoeflich et al., 2004). However, BMPR-2 lacking this region (*Bmpr2Δ751-813aa*) rescued the phenotype of *Limk1* overexpression. This suggest that BMPR-2 inhibits LIMK with the tail domain but not at 751-813aa at least in mitral cells. Age, P6. Arrows and arrowheads indicate somata and primary dendrites, respectively. Scale bar, 100 μm. **(C)** Quantification of the number of glomeruli innervated per mitral cell. n, number of mitral cells. n.s., non-significant (χ^2^ test, compared to the control).

**Figure S3.**
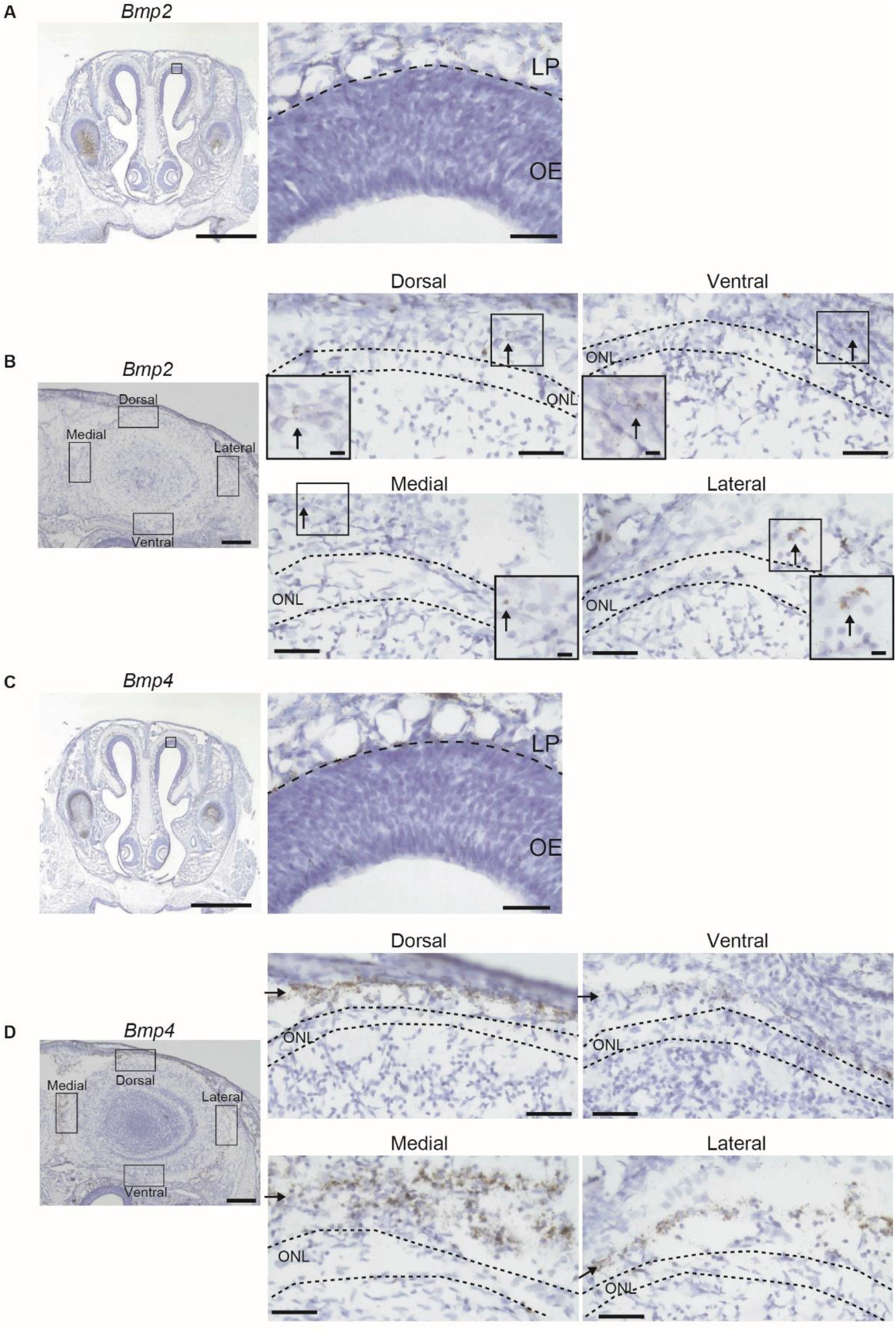
Expression of *Bmp2* and *Bmp4*. **(A, B)** Expression of *Bmp2* visualized by RNA scope combined with DAB staining. *Bmp2* expression was observed in the lamina propria (LP) but not in the OE. *Bmp2* was also expressed on the OB surface, but at a lower level. Arrows indicate DAB signals. Insets show high magnification images of areas indicated by arrows. **(C, D)** *Bmp4* expression was observed in the LP but not in the OE. *Bmp4* was also expressed on the OB surface, although the expression level was lower in the ventral part. Arrows indicate DAB signals. Scale bars are 1 mm (A, C, left), 250 μm (B, D, left), 50 μm (middle and right), and 10 μm (insets).

**Figure S4.**
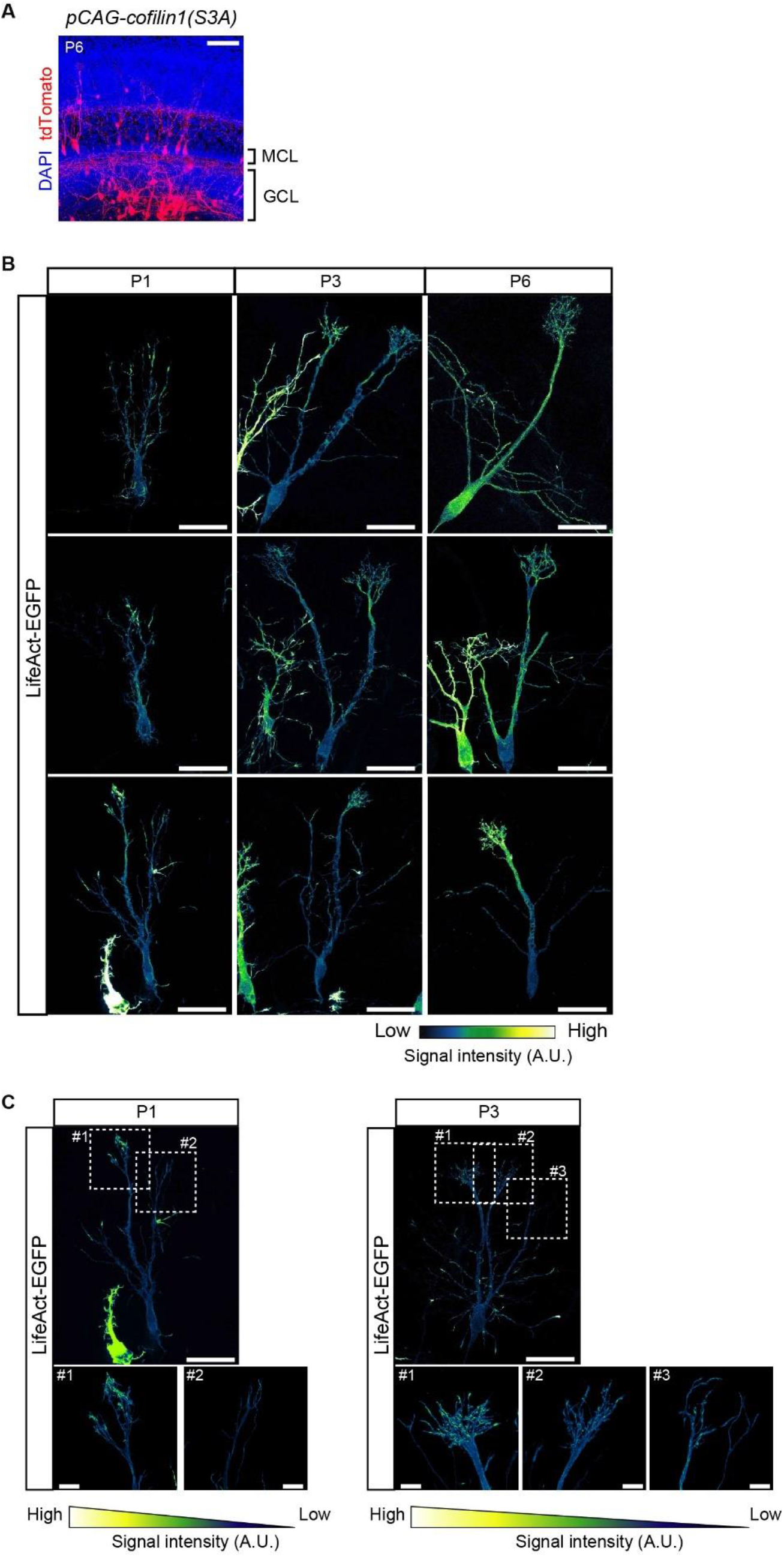
Localization of F-actin. **(A)** Overexpression of *cofilin1(S3A)* under a constitutively active CAG promoter. Many mitral cells were stacked in the granule cell layer (GCL), most likely due to a migration defect. MCL, mitral cell layer. Scale bar, 100 μm. **(B)** Additional examples of LifeAct-EGFP localization at different stages of dendrite remodeling. The signal intensity was adjusted to the brightest signal at the dendritic tufts. Scale bars are 50 μm. **(C)** In some mitral cells with multiple tufted dendrites, LifeAct signals were biased to one of multiple dendrites extending to the glomerular layer. In these examples, one of the dendritic tufts (#1) were brighter than the others. Here, the dendrite #1 had more developed tufted structure. This may indicate that F-actin formation is promoted in a prospective winner dendrite.

**Figure S5.**
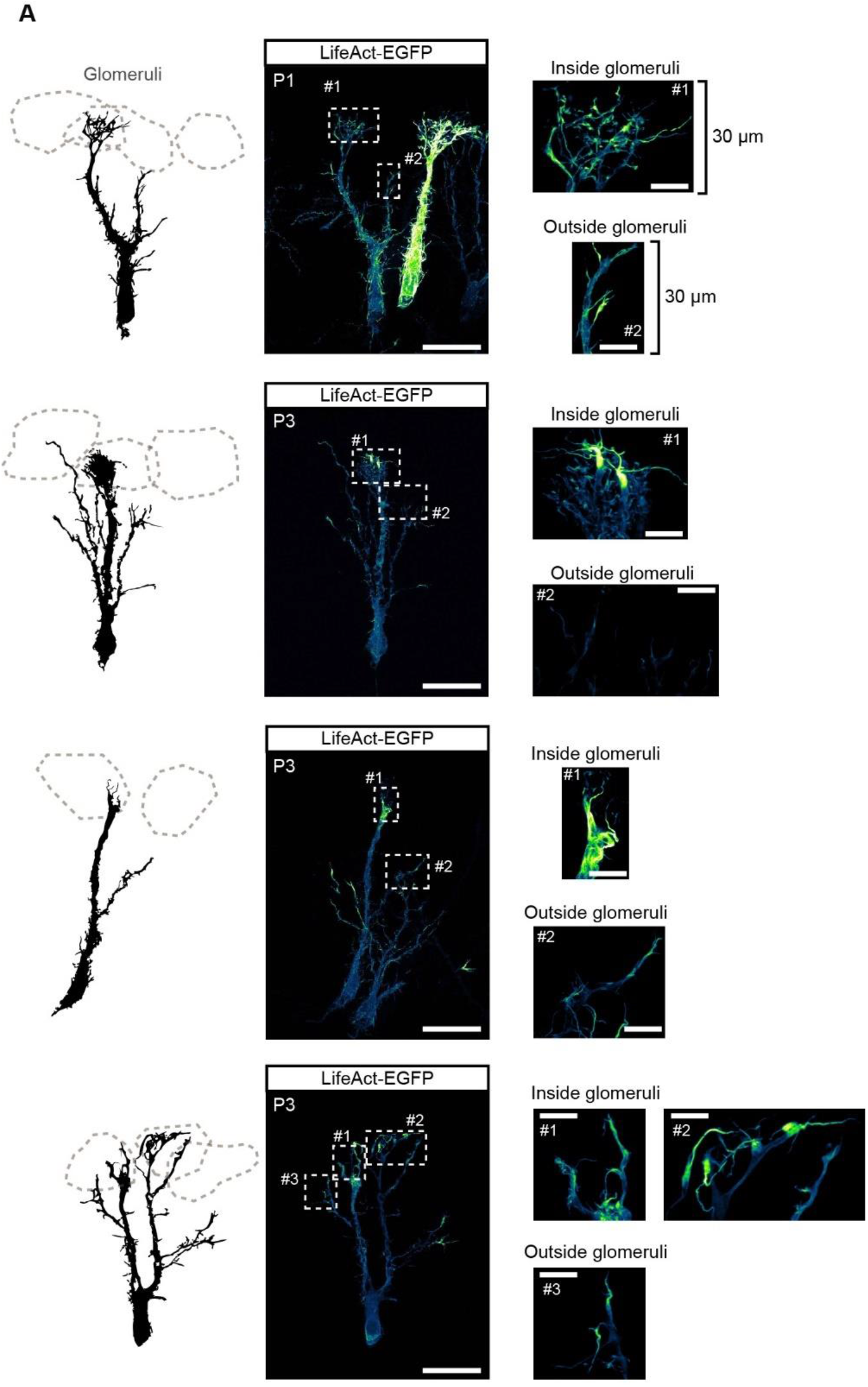
Comparison of LifeAct signals between dendrites inside and outside glomeruli. **(A)** Additional images of mitral cells extending dendrites inside and outside glomeruli. Traces are shown on the left. LifeAct signals were most prominent in dendritic terminals inside glomeruli. Age, P1 or P3. Scale bars are 50 μm (left) and 10 μm (right).

